# Sex Based Differences in Active Avoidance and Approach Strategy in the Platform Mediated Avoidance Task

**DOI:** 10.1101/2025.10.01.679640

**Authors:** Christina J. Li, Delilah Pineda, Adriano E. Reimer, Steven M. Hu, Michael R. Angstman, Jason L. Chang, Alik S. Widge

**Affiliations:** Department of Psychiatry & Behavioral Sciences, University of Minnesota, Minneapolis, MN, USA

**Author notes:** Corresponding Author: Alik S. Widge, Department of Psychiatry & Behavioral Sciences, University of Minnesota, Minneapolis, MN 55455. These authors contributed equally to this work (Christina J. Li, Delilah Pineda, Adriano E. Reimer).

**Keywords:** Sex Differences, Approach-Avoidance Conflict, Platform Mediated Avoidance (PMA), Active Avoidance, Anxiety Disorders

## Abstract

Sex differences have been documented in threat conditioning, but research into potential sex differences in avoidance paradigms, particularly active avoidance, remains limited. This research gap is particularly concerning given that women are disproportionately affected by stress- and anxiety-related disorders, characterized by maladaptive avoidance. Yet, preclinical research has historically focused on male subjects, limiting our understanding of the neurobiological mechanisms underlying sex differences in threat responses. To address this, we investigated sex-specific strategies in adult Long Evans rats (10 female, 9 male) using a modified platform-mediated avoidance (PMA) task that created a high-conflict choice between reward-seeking and safety. Behavior was tracked over 25 days, with analyses focusing on a stable performance phase (days 20-25) objectively defined using change point analysis. The study design included an initial cohort and a replication cohort to ensure the findings’ robustness. Females consistently prioritized safety, spending significantly more time foregoing reward to avoid foot shock, while males engaged in more persistent reward-seeking despite the risk of shock. This difference was not driven by differential reward motivation. Furthermore, female strategies were not significantly modulated by the estrous cycle. Thus, male and female rats employ fundamentally different strategies to resolve approach-avoidance conflict: females adopt a robust, safety-first strategy, while males demonstrate a risk-prone, reward-oriented approach. These findings highlight the importance of considering biological factors underlying threat responses, suggesting that characterizing these neural mechanisms may guide more targeted interventions for anxiety and trauma-related disorders.

## Introduction

Women are substantially more likely than men to be diagnosed with stress- and anxiety-related disorders, including post-traumatic stress disorder (PTSD) and generalized anxiety disorder (GAD) (Kessler et al., 2005; McLean et al., 2011, Haering et al., 2024). These conditions are highly comorbid, share overlapping symptoms (Price & van Stolk-Cooke, 2015; Afzali et al., 2017), and rank among the most prevalent mental health challenges globally, contributing to significant societal and economic costs (Fineberg et al., 2013; Trautmann et al., 2016). Despite the clear sex disparity in prevalence, the underlying neurobiological mechanisms driving these differences remain poorly understood (Lebron-Milad & Milad, 2012; Day & Stevenson, 2020).

A core behavioral symptom and transdiagnostic feature across these disorders is maladaptive avoidance, where individuals excessively avoid situations or stimuli perceived as threatening (Dymond, 2019). Frequently arising from attempts to regulate intense negative emotions, this strategy involves preventing contact with sources of distress by including avoidance of external triggers and the avoidance of internal thoughts and feelings (Barlow, 2008; Ellard et al., 2010). Because this pattern can severely impair daily functioning, it is a key target for clinical intervention (Salters-Pedneault et al., 2004; Blakey & Abramowitz, 2016). Examining the basis of sex differences in the expression of avoidance, which allows for better understanding of the dysfunctional mechanisms underlying the mental disorders, is therefore a critical step toward developing more effective, personalized treatments.

Rodent models have been instrumental in elucidating the neural circuits of threat conditioning and avoidance (LeDoux, 2000; Maren, 2001; Bienvenu et al., 2021; Diehl et al., 2024), yet preclinical research has historically focused almost exclusively on male subjects (Beery & Zucker, 2011; Dalla et al., 2009; Shansky, 2018). This persistent bias has limited our ability to model the full spectrum of anxiety pathology and has created significant gaps in our knowledge of the female neurobehavioral response to threat (Shansky, 2019; Shansky & Murphy, 2021).

The limited studies that have included both sexes reveal important sexual dimorphisms. For instance, female mice can demonstrate more persistent avoidance following extinction training (Halcomb et al., 2024), and both female rats and mice often employ different defensive response strategies compared to males, such as active darting or “anxioescapic” behaviors instead of passive freezing (Gruene et al., 2015; Shanazz et al., 2022; Halcomb et al., 2024). This divergence may reflect fundamentally different cognitive strategies for assessing risk. For instance, in value-based decision-making, females are more likely to use a steady, long-term “less risky” strategy, whereas males often change their approach based on recent outcomes (Chen et al., 2021a, 2021b). Such findings suggest that sex-based variations could arise from diverse cognitive approaches (Grissom et al., 2024). The hormonal contribution to these different approaches, however, is not straightforward; for example, although hormonal fluctuations across the estrous cycle in females can modulate some behaviors (Reimer et al., 2018; Maeng et al., 2017), its influence is often limited or absent in others (Zeng et al., 2023; Levy et. al., 2023; Becegato & Silva, 2024).

While identifying these distinct strategies, and the underlying mechanisms, is a crucial step, the clinical relevance of these findings may be limited by paradigms that do not fully capture this strategic complexity (Aupperle & Paulus, 2010; Kirlic et al., 2017, Ball & Gunaydin, 2022). Avoidance tasks are typically categorized as passive (withholding an action) or active (performing one). Focusing on the latter, many standard active avoidance paradigms, like the shuttle box, have translational limitations (Diehl et al., 2019). The shuttle avoidance (Mowrer & Lamoreaux, 1946; Krypotos et al., 2015), for example, requires an animal to repeatedly re-enter a compartment where it was just shocked. This creates a condition where there is no well-defined safe location, leading to an ambiguous response: it promotes freezing behavior that directly competes with, and can be mistaken for, the avoidance strategy being measured. Furthermore, these tasks typically omit any cost for the avoidant action, failing to model the clinically relevant situations where individuals sacrifice valuable opportunities by engaging in excessive, maladaptive avoidance that severely impairs their daily functioning (Aupperle et al., 2015; Pittig et al., 2021).

More recently, the Platform-Mediated Avoidance (PMA; Bravo-Rivera et al., 2014) task was developed to address these issues, allowing for a less ambiguous behavioral response by providing a distinct safe zone while also incorporating a cost-benefit component. However, this paradigm models a specific type of conflict where the threat and reward are separated in time; an animal can simply wait for the threat to pass before safely resuming reward-seeking. While this is a valuable model, it does not fully capture the decision-making characteristic of clinical anxiety, where valuable opportunities are contingent upon enduring a perceived threat. A more direct model requires forcing an explicit trade-off where the reward is only available in the presence of the threat. Investigating the potentially dimorphic strategies that emerge under such intense conditions - which can reflect differences in threat-response regulation, decision-making, or both (Orsini et al., 2016; Greiner et al., 2019; Xu et al., 2022) - is a crucial step toward understanding the neurobiological basis of avoidance circuits and how they may be altered in clinical populations (Olff, 2017; Panayiotou et al., 2017).

Therefore, the present study employed a modified PMA task designed to create the type of intense approach-avoidance conflict needed to reveal these strategies. In our paradigm, animals could only access rewards when a threat was imminent, forcing a choice between reward-seeking and safety. We tracked the development of avoidance and approach behaviors over 25 days to examine how strategies were acquired and expressed once they stabilized, allowing us to compare both the learning dynamics and end-stage behavioral patterns between sexes.

Based on the clinical prevalence of anxiety disorders in women and preclinical evidence of dimorphic threat responses, we hypothesized that male and female rats would adopt distinct strategies. Specifically, we predicted that females would exhibit greater avoidance behavior, prioritizing safety at the cost of reward, whereas males would demonstrate more persistent reward-seeking. We also conducted exploratory analyses to determine whether behavioral strategies in females were modulated by the estrous cycle, given the potential influence of reproductive hormones on threat-detection / threat-response in rodents.

## Methods

### Preregistration

The first cohort (4 female, 4 male) was not preregistered. To test whether the exploratory findings from that cohort were replicable, the experimental design and analysis plan for the second cohort (6 female, 5 male) were preregistered on the Open Science Framework (https://osf.io/gav3h) prior to data collection. The sample size was determined via power analysis for repeated measures ANOVA assuming a small effect size (f^2^ = 0.1), alpha of 0.05, and power of 0.80 (full details in preregistration), used as a conservative approximation for the planned GLMMs. This analysis indicated that N = 10 was sufficient to detect small-to-moderate effects.

### Animals

All experiments were conducted in accordance with the NIH Guidelines for the Care and Use of Laboratory Animals and were approved by the University of Minnesota Institutional Animal Care and Use Committee (IACUC, protocol 2303-40918A). A total of 19 adult Long Evans rats (10 female, ∼250g; 9 male, ∼350g; ∼3 months old), purchased from Charles River Laboratories (Wilmington, MA), were used in this study. Upon arrival, rats were pair-housed under a 12-hour reverse light/dark cycle (lights off at 9:00 AM) and allowed to acclimate for at least 7 days. Following acclimation, they were individually housed and handled daily for 7 consecutive days. To maintain motivation for the sucrose reward, rats were food restricted to 90% of their baseline body weight. Their weight was monitored daily, and food allotment was adjusted to maintain this target weight (not falling below 85% of baseline). All behavioral testing occurred during the dark phase under red light.

Animals were tested in two cohorts. The first cohort (4F, 4M) was run prior to preregistration. The second, preregistered cohort (6F, 5M) served as a replication. Behavioral modeling was conducted separately for each cohort to preserve the confirmatory nature of the preregistered analyses. Data from both cohorts were combined for estrous-stage analyses, which were designated as exploratory in the preregistration, to increase statistical power.

### Apparatus

Behavioral testing was conducted in modified operant chambers (40.6 cm × 25.4 cm × 44.0 cm; Lafayette Instruments, Lafayette, IN) housed within sound-attenuating cubicles. Each chamber featured a grid floor capable of delivering scrambled footshocks, a retractable lever with a cue light and speaker, and pellet dispenser and trough on one wall, and an acrylic safety platform (10.2 cm × 25.4 cm) on the opposite wall. Sessions were recorded by digital video cameras (DMK 37AUX287, The Imaging Source, NC) mounted above the behavioral chamber. A computer and interface unit (Lafayette Instruments) was connected to the chambers, and all task parameters, stimulus presentation, and data acquisition were controlled by in-house developed behavior control software Pybehave (Dastin-van Rijn et al., 2024).

### Procedures

#### Platform Mediated Avoidance Task

The behavioral protocol consisted of three sequential stages: (1) initial lever press training, (2) conditioning of separate threat and reward cues, and (3) an extended high-conflict PMA task (Figure 1-A).

**Figure 1.**
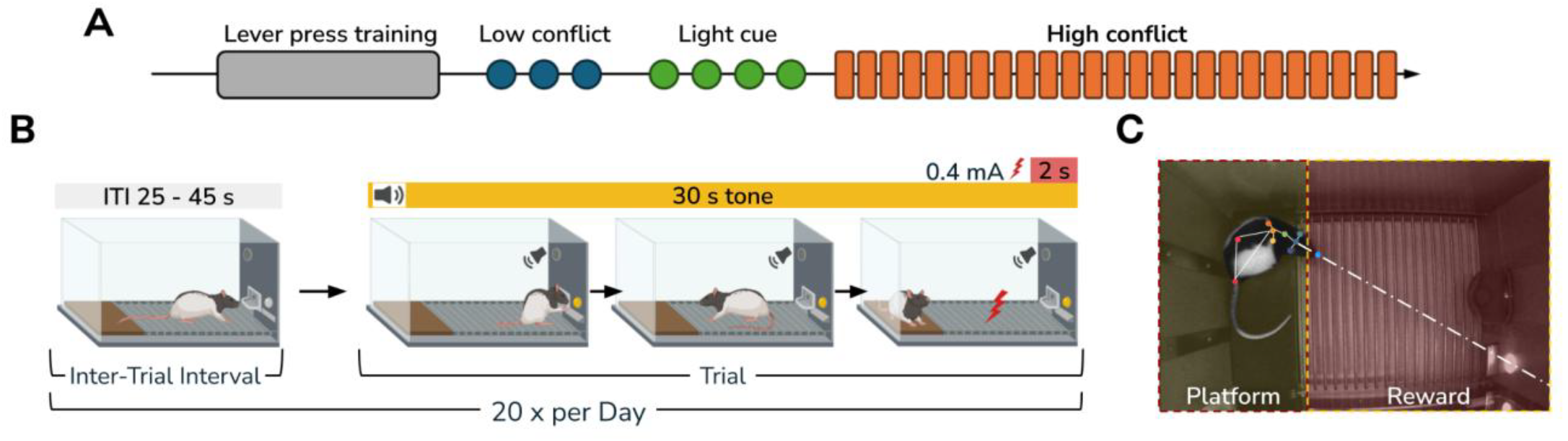
Experimental timeline and Platform Mediated Avoidance (PMA) task paradigm. (A) Experimental timeline: After habituation and once target weight was reached through food restriction, rats were trained to lever press for a food reward. This was followed by low conflict (3 days), light cue association (4 days), and high conflict (25 days) sessions. (B) High conflict PMA task: Each daily session consists of 20 trials (variable inter-trial interval, ITI, 25-45 seconds). A trial begins with the onset of a 30-second auditory tone. Only during this tone, the rat can press the lever to receive a food reward but must retreat to the safe platform to avoid a footshock (0.4 mA for 2 seconds) delivered through the grid floor at the end of the tone. (C) Still frame showing tracked body points using DeepLabCut. Zones of interest (platform and reward, shaded in yellow and red, respectively). The frame also illustrates the assessment of reward attentiveness, where the animal’s head is oriented toward the reward zone (indicated by the eyeline vector).

First, all animals underwent operant conditioning to press a lever for 45-mg sucrose pellets in 40-minute daily sessions. Training began on a fixed-ratio 1 (FR1) schedule and advanced to a variable-interval 30-second (VI30s) schedule after an animal achieved a rate of at least 10 presses within the first 10 minutes of a session. The same criterion was required on the VI30s schedule before proceeding.

After lever-press acquisition, rats underwent two distinct pre-training phases to establish key associations. First, in a three-day, low-conflict conditioning phase, a 6 kHz warning tone was paired with a co-terminating 2-second, 0.4 mA footshock over twenty 30s trials (Inter-Trial-Interval, ITI: 25-45s). The lever remained continuously available during the ITI. Next, animals were subjected to a four-day reward-cue phase, in which the lever and an associated cue light were made accessible only during the 30 s trial periods, with no tones or shocks delivered. This trained the animals to associate the light-cued trial period with reward availability.

The final phase was a 25-day high-conflict PMA task. The trial structure (twenty 30s trials; 25-45s ITI) was maintained. Critically, the food lever and its light cue were now accessible only during the presentation of the tone, which still co-terminated with the footshock. This design created a direct approach-avoidance conflict, forcing animals to choose between approaching the lever for a reward and retreating to the platform for safety (Figure 1-B). At the end of each trial, the lever was retracted, becoming unavailable during the ITI.

#### Estrous Cycle Staging

Following each behavioral session, vaginal lavage was conducted on female rats to determine estrous stage. It was performed using a pipette containing approximately 100 µL of sterile saline. Saline was gently flushed in and out of the vaginal canal several times to collect epithelial cells. The sample was then transferred to a glass slide. The slides were left to dry in room temperature followed by staining (DipQuick Stain Fixative, Stain Solution, and Counter Stain, JorVet, CO) and examination under a light microscope for cytological assessment (Cora et al., 2015). To control for stress associated with this procedure, male rats were held in the same restraint position for a similar duration after each session.

### Behavioral and Statistical analysis

#### Pose Estimation

Animal body position was tracked using DeepLabCut (v3.0RC10; Mathis et al., 2018, Nath et al., 2019). A ResNet-50-based neural network (Insafutdinov et al. 2016; He et al. 2016) with default parameters was used on 1230 manually labeled frames from 108 video fragments including different animals/sessions (95% training, 5% testing), for 6 shuffles. The final trained network achieved a test error of 2.61 pixels and a training error of 1.6 pixels (image size: 720 × 540 pixels) with a p-cutoff of 0.6.

#### Post-Processing

All subsequent behavioral data processing was performed in the RStudio environment (v2025.5.1.513; Posit team, 2025) using R (v4.5.1; R Core Team, 2025). Data wrangling and transformation were conducted primarily using the tidyverse (Wickham, 2023; v2.0.0), including its component packages dplyr (Wickham et al., 2023; v1.1.4), tidyr (Wickham & Girlich, 2024; v1.3.1), tibble (Müller & Wickham, 2025; v3.3.0), stringr (Wickham, 2025; v1.5.1), purrr (Henry & Wickham, 2025; v1.1.0), readr (Wickham et al., 2024; v2.1.5), and glue (Hester, 2024; v1.8.0). In addition, data.table (Dowle & Srinivasan, 2025; v1.17.8) was used for data aggregation.

The behavioral chamber was divided into two zones: platform and reward (Figure 1-C). Events (tone, shock, lever press) were extracted from PyBehave log files. An animal’s location was determined by the coordinates of its lower back. Time spent on the platform was defined as the duration the lower back was within the platform zone. Time in the reward zone was defined similarly. Reward attentiveness was calculated as the total time an animal spent either physically within the reward zone or oriented toward it from outside the zone. Head orientation was determined by calculating a forward-projecting vector from the animal’s head-center through its nose on a by-frame basis. If this vector intersected with the reward zone’s coordinates, the frame was counted as “looking at the reward.”

#### Change Point Analysis

To objectively identify the onset of stable, end-stage behavior, Bayesian change point analysis was applied to the daily averages of three key metrics from Cohort 1: percent time on the platform, reward attentiveness, and total bar presses. Using the mcp package in R (Lindeløv, 2024; v0.3.4), performance for each metric and sex was modeled with a three-segment trajectory: 1) a flat baseline, 2) a quadratic learning phase, and 3) a final flat stable phase (model: DailyAvg ∼ 1, ∼ 0 + DAY + I(DAY^2), ∼ 1). This extended learning phase is characteristic of the PMA paradigm, which requires a prolonged period for the initial freezing response to subside and for competing behaviors, such as reward-seeking, to stabilize (Bravo-Rivera et. al., 2014; Martínez-Rivera et. al., 2020).The second change point, marking the transition into the stable phase, was estimated for each model. The latest of these change points across all metrics and both sexes was Day 19. Therefore, performance on days 20–25 was defined as the stable phase for both cohorts.

#### Behavioral Modeling

Stable-phase behavior was analyzed using generalized linear mixed models (GLMMs) in R, with models fit separately for each cohort (Bolker et al., 2009). The high-conflict design of the task, forcing a choice between the safety of the platform and the reward at the lever, produced bimodal data distributions. Specifically, many trials resulted in mutually exclusive behavioral outcomes, where an animal either remained on the platform for the entire duration (resulting in 100% platform time and zero bar presses) or committed to reward-seeking by approaching the lever. To properly capture this structure, we selected two-part models that separate the binary choice to engage in a behavior from the subsequent frequency or duration of that behavior (Fisher et al., 2017; Zhu et. al., 2017). Based on data characteristics from Cohort 1, three primary dependent variables were modeled:

1. Bar Presses: Modeled using a zero-inflated negative binomial (ZINB) GLMM. The model’s zero-inflation component estimated the probability of a rat not approaching the lever at all within a trial, while the negative binomial component modeled the number of bar presses on trials where it did approach.
2. Proportion of Time on Platform: Modeled using a hurdle-beta GLMM. A binomial model first predicted the probability of spending any time off the platform (hurdle), followed by a beta regression on the proportion of time for trials where the animal did not spend 100% of the time on the platform.
3. Reward Attentiveness: Modeled using a beta GLMM to analyze the proportion of trial time the animal was oriented toward the reward zone.

For both proportion-based measures (platform time and attentiveness), values were transformed using the Smithson and Verkuilen (2006) method to fit the open (0,1) interval required for beta regression.

Each model included fixed effects for sex (female = 0, male = 1), day (continuous), trial (1-20, continuous), handler (categorical), previous shock history on that day (SH; binary). To account for repeated measures, animal ID was included as a random intercept. An important consideration for the analysis was the inherent correlation between sex and body weight, as adult male and female rats have distinct weight distributions. To create a predictor for relative size that was independent of sex, we transformed each animal’s absolute weight by z-scoring it against the mean and standard deviation of all weights from its respective sex, calculated across the stable phase (days 20-25). This average z-scored weight was then included as a fixed effect in the models. The preregistered analysis plan also included an interaction terms for sex-by-weight. However, post-hoc multicollinearity diagnostics revealed that the sex-by-weight interaction introduced statistical instability, indicated by Variance Inflation Factors (VIFs) exceeding 5 (Kim, 2019). Therefore, in a deviation from the preregistration, this term was removed from the final reported models to ensure their validity and the reliability of the coefficients.

All continuous predictors were normalized to a 0-1 scale. Statistical significance was set at p < 0.05. Models were fit using the glmmTMB package (Brooks et al., 2017; v1.1.11) and lme4 package (Bates et al., 2015; v1.1.37), with alternative optimization algorithms accessed via optimx (Nash & Varadhan, 2025; v2025-4.9) when convergence issues arose. Residual and model diagnostics were performed using DHARMa (Hartig, 2024; v0.4.7), performance (Lüdecke et al., 2025; v0.15.0), and see (Lüdecke et al., 2025; v0.11.0). Robust variance estimates were checked using sandwich (Zeileis et al., 2024; v3.1.1) and lmtest (Hothorn & Zeileis, 2022; v0.9.40).

#### Estrous Based Analysis

To explore hormonal influences on stable behavior, we conducted an exploratory analysis using trial-level data from female rats (n = 10) across both cohorts during the stable phase (Days 20-25). Daily cytological classifications were grouped into a binary “Hormone_Level” factor: “High-Hormone” (proestrus, estrus) and “Low-Hormone” (metestrus, diestrus), following Reimer et al. (2018), with the latter as reference category. We applied the same GLMM framework used in the primary analysis, with Hormone_Level replacing Sex as the primary predictor of interest. The Reward Attentiveness (beta GLMM) and Platform Time (hurdle-beta GLMM) models successfully converged with all preregistered covariates. However, the Bar Presses (ZINB) model initially failed to converge due to statistical instability from the reduced female-only sample size. To achieve a stable model while preserving the most influential variables, we removed the average z-scored weight term from the conditional component, as this was the least consistent predictor across cohorts in the main analysis. The zero-inflation component retained all covariates. All models included animal ID as a random intercept.

#### Visualization and reporting

Figures were generated using ggplot2 (Wickham et al., 2025; v3.5.2), with palettes from RColorBrewer (Neuwirth, 2022; v1.1.3) and additional layers from ggnewscale (Campitelli, 2025; v0.5.2) and ggsignif (Ahlmann-Eltze & Patil, 2022; v0.6.4). Plot layouts were composed with patchwork (Pedersen, 2025; v1.3.1). Tables were formatted with gt (Iannone et al., 2025; v1.0.0). Model outputs were tidied for reporting using broom.mixed (Bolker & Robinson, 2024; v0.2.9.6).

## Results

### Change point analysis

To objectively define the transition from learning to stable behavior, we conducted a Bayesian change point analysis on the daily performance metrics of the initial cohort (Cohort 1). We fit a three-segment piecewise regression model (flat baseline, quadratic learning, flat stable phase) to three behaviors: avoidance, reward attentiveness, and bar presses. Models were run for 700,000 iterations across 7 chains to ensure convergence. For females, the transition to the stable plateau occurred at Day 11.36 for avoidance (95% CI: 8.00-14.96), Day 12.46 for reward attentiveness (95% CI: 6.20-25.00), and Day 13.07 for bar presses (95% CI: 9.44-17.00). Males exhibited a more extended learning phase, with behavior stabilizing at Day 16.70 for avoidance (95% CI: 13.00-22.57), Day 14.74 for reward attentiveness (95% CI: 8.00-24.03), and Day 18.92 for bar presses (95% CI: 14.71-23.00). To establish a conservative window ensuring all animals had reached a performance asymptote, we defined the stable phase based on the latest change point. With the final transition occurring at Day 19 for male bar pressing, all subsequent analyses were conducted on data from experimental days 20-25 (Figure 2).

**Figure 2.**
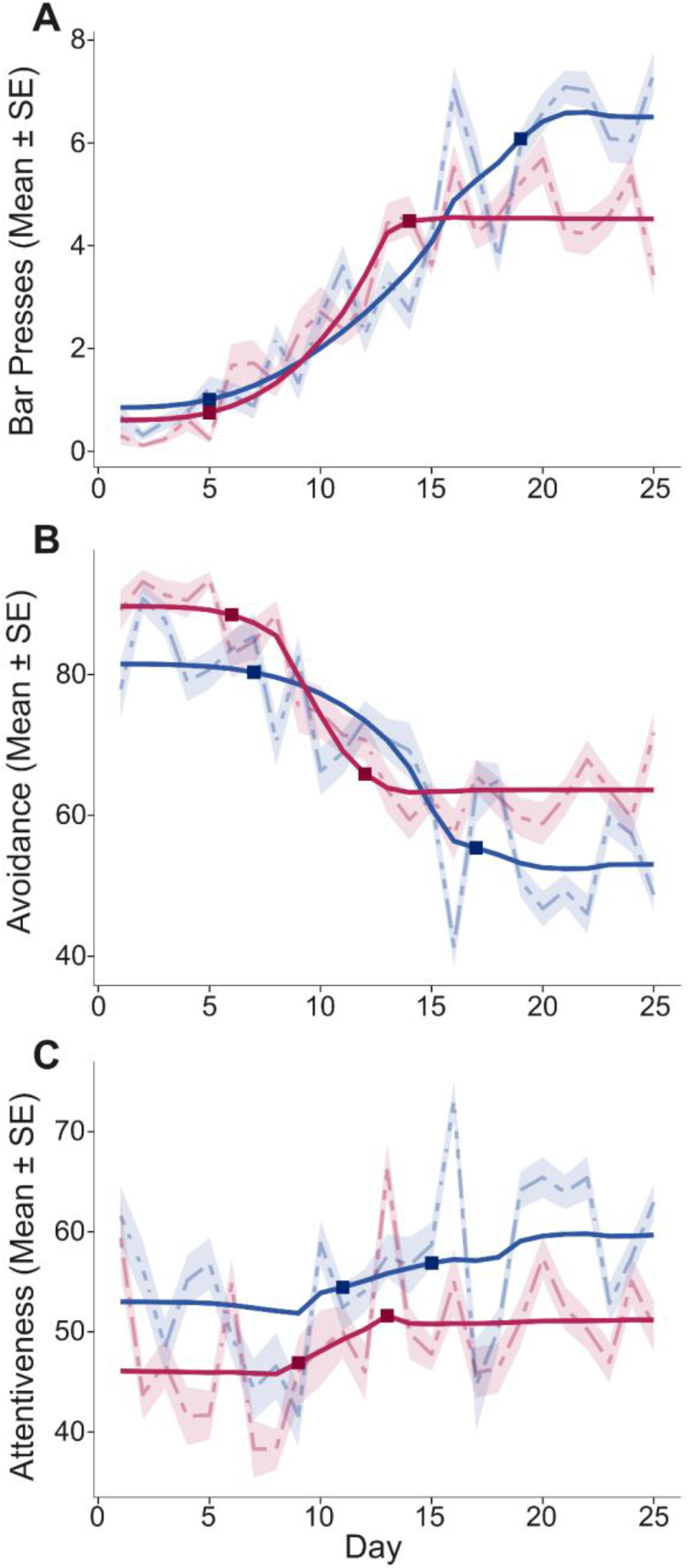
Determination of Stable Behavior Using Change Point Analysis. The onset of stable performance in Cohort 1 (n=8; 4 females, 4 males) was modeled across three key behaviors: (A) total daily bar presses, (B) percent of time spent on the platform, and (C) percent of time attentive to the reward zone. Dotted lines represent observed means ± standard error; solid lines show the mean posterior prediction of the fitted segmented regression model. Square markers denote estimated change points (τ).

### Initial study and replication

To investigate sex differences in avoidance and reward-seeking strategies during the stable phase of behavior (days 20-25), we analyzed three key metrics: bar presses, avoidance (proportion of time on the platform), and reward attentiveness. GLMMs were fit separately for Cohort 1 and Cohort 2, as per the preregistered plan.

### Cohort 1

#### Reward-Seeking Behavior (Bar Presses)

Analysis with the ZINB model showed non-significant increase in male reward seeking compared to females (β = 0.15, z = 1.83, p = 0.067; Figure 3-A). SH was a significant negative predictor of bar pressing (β = -0.33, z = -7.97, p < 0.001). Additionally, the animal’s average z-scored weight was a significant positive predictor of bar presses (β = 0.26, z = 2.83, p = 0.005). The zero-inflation portion of the model, which predicts the likelihood of an animal making no presses, showed that both SH (β = 0.66, z = 2.84, p = 0.005) and day (β = 0.93, z = 2.60, p = 0.009) significantly increased the probability of complete inaction in a trial.

**Figure 3.**
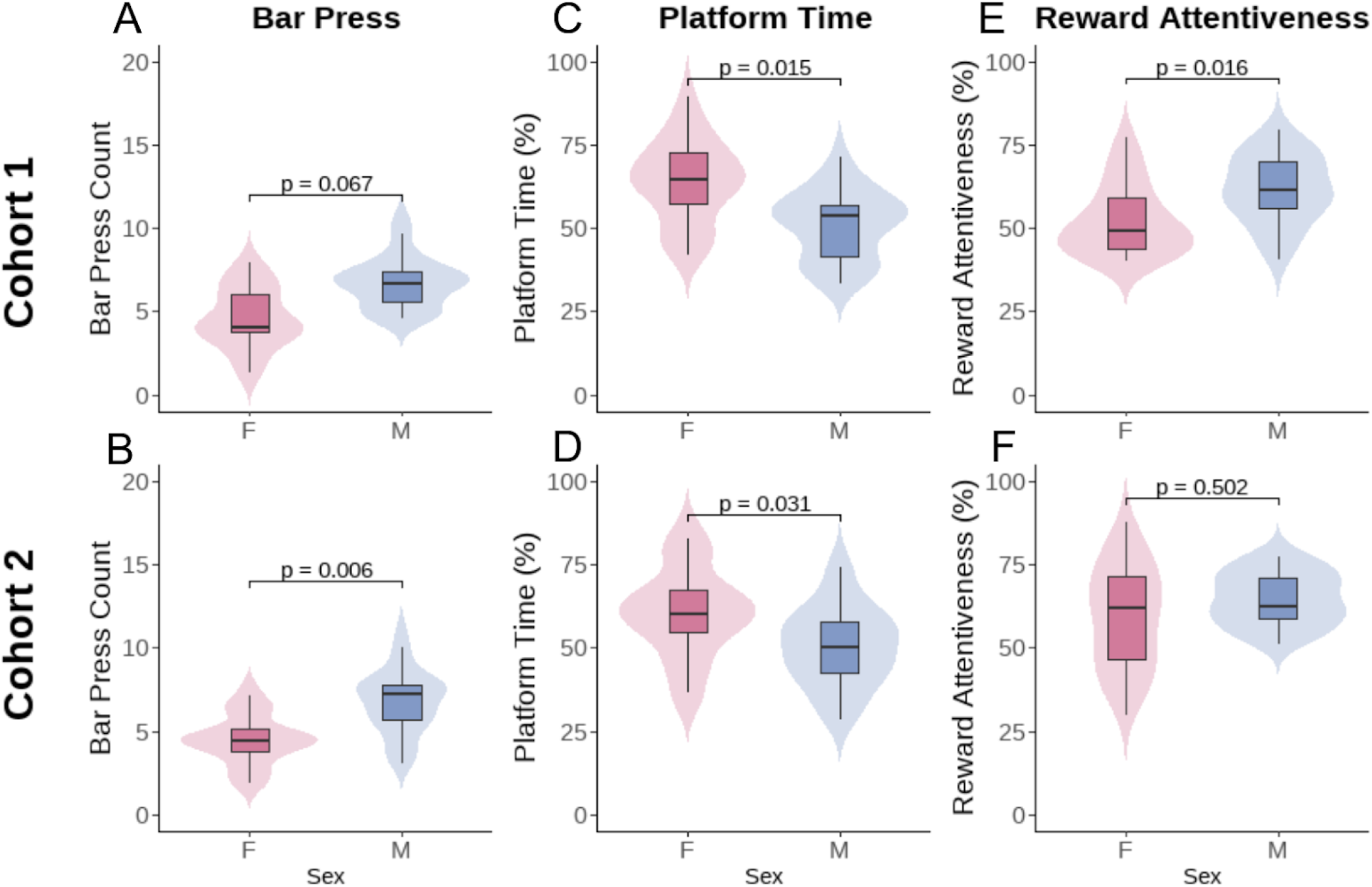
Sex differences in stable-phase performance in the high conflict task. Performance during Days 20-25 is shown for Cohort 1 (top row; n = 4F, 4M) and the replication Cohort 2 (bottom row; n = 6F, 5M). Violin plots with overlaid boxplots comparing females and males across three metrics: (A,B) Bar Press Count, (C,D) Platform Time (%), and (E,F) Reward Attentiveness (%). P-values from statistical comparisons between sexes are displayed above groups.

#### Avoidance Behavior (Platform Time)

Avoidance behavior was analyzed with a two-part hurdle-beta model. The first part (binomial hurdle) revealed no sex difference in the decision to leave the platform at any point during a trial (p = 0.124). However, the second part of the model (beta), analyzing the trials where animals actually left the platform, revealed a significant main effect of sex, with males spending less time on the platform than females (β = -0.30, z = -2.44, p = 0.015; Figure 3-C). Avoidance in this cohort was also significantly predicted by SH (β = 0.68, z = 9.43, p < 0.001), trial (β = -0.33, z = -3.22, p = 0.001), and weight (β = -0.54, z = -4.33, p < 0.001).

#### Reward Attentiveness

The beta GLMM for reward attentiveness revealed a significant main effect of sex. Males were significantly more attentive to the reward zone than females (β = 0.42, z = 2.40, p = 0.016; Figure 3-E). SH also significantly decreased reward attentiveness (β = -0.27, z = -4.56, p < 0.001).

We then pre-registered these findings as predictions for a replication analysis, powered to detect the effect sizes observed in Cohort 1.

### Cohort 2

#### Reward-Seeking Behavior

The ZINB model revealed a significant main effect of sex, with males exhibiting a higher rate of bar pressing than females (β = 0.44, z = 2.74, p = 0.006; Figure 3-B), reinforcing the trend observed in Cohort 1. Similar to the first cohort, SH significantly reduced bar pressing (β = - 0.15, z = -4.59, p < 0.001), while both handling groups H2 and H3 showed increased pressing compared to the reference group. The model also revealed a significant sex-day interaction (β = -0.16, z = -1.97, p = 0.049) which was not observed in Cohort 1. The zero-inflation part of the model also replicated the first cohort’s results, showing that SH significantly increased the probability of a complete lack of reward-seeking on the subsequent trial (β = 0.73, z = 3.08, p = 0.002).

#### Avoidance Behavior

The hurdle model again revealed no significant sex difference in the decision to leave the platform (p = 0.888). However, SH was a significant predictor (β = 0.66, z = 2.28, p = 0.023). Critically, the second part of the model (beta regression) confirmed the significant main effect of sex, (β = -0.60, z = -2.16, p = 0.031; Figure 3-D). Avoidance was also strongly predicted by SH (β = 0.47, z = 7.59, p < 0.001), day (β = -0.40, z = -4.03, p < 0.001), and trial number (β = -0.44, z = -5.14, p < 0.001). A significant sex-day interaction was also found (β = 0.38, z = 2.63, p = 0.008).

#### Reward Attentiveness

In contrast to the first cohort where a significant sex difference was found, the beta GLMM for reward attentiveness in the replication cohort did not show a significant main effect of sex (p = 0.502; Figure 3-E). However, consistent with Cohort 1, attentiveness was significantly reduced by SH (β = -0.50, z = -9.33, p < 0.001).

The cross-cohort comparisons of our findings is summarized in Figures 4, 5 and 6, which compare the model predictors for bar pressing, avoidance behavior, and reward attentiveness between cohorts.

**Figure 4.**
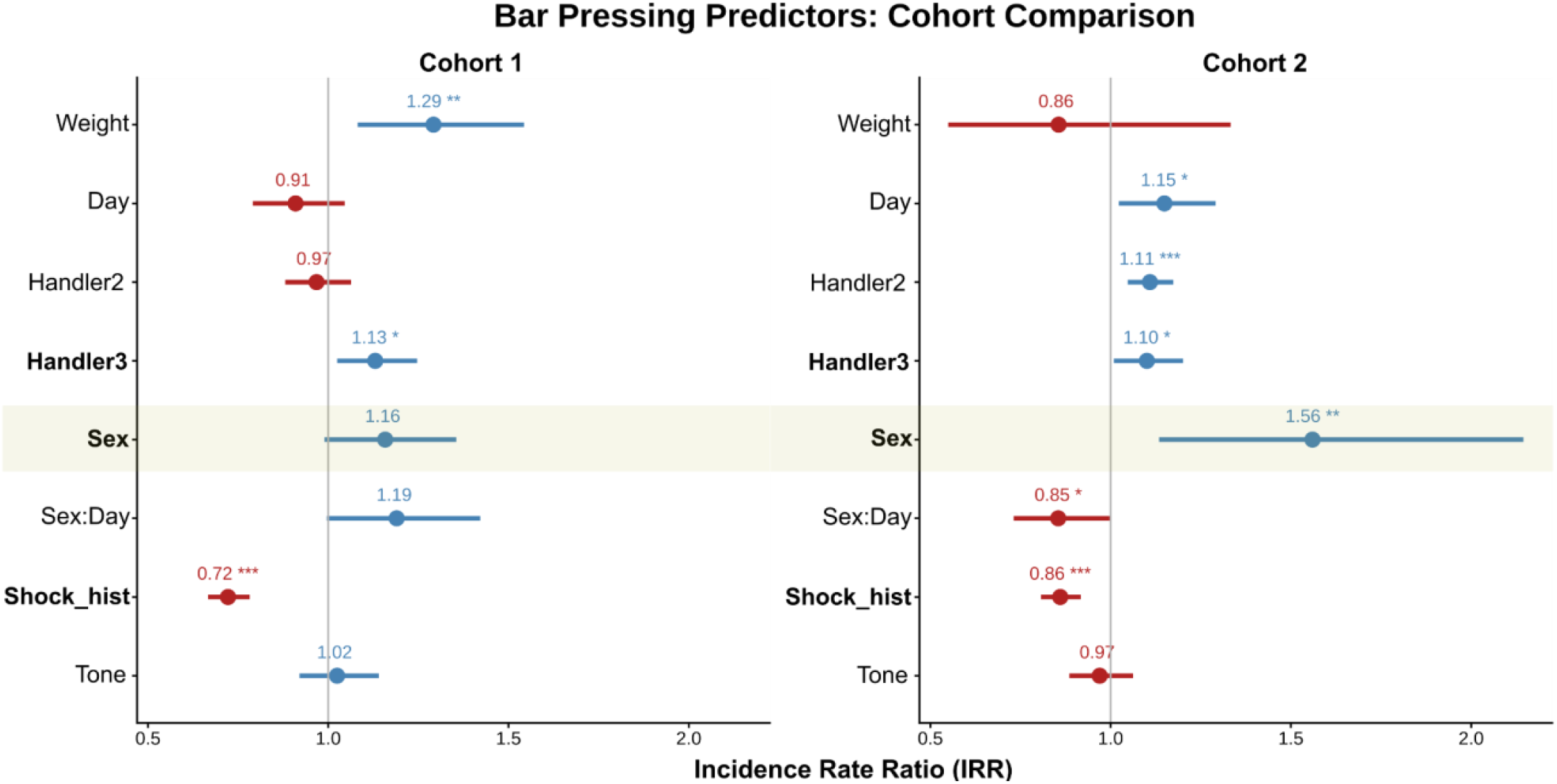
Comparison of predictors for bar pressing across cohorts. Forest plots display Incidence Rate Ratios (IRRs) with 95% CI from the GLMMs for Cohort 1 (left) and Cohort 2 (right). IRRs represent the multiplicative effect of each predictor on the rate of bar pressing; values > 1 (blue) indicate a positive association, while values < 1 (red) indicate a negative one. Predictors include: Sex (male vs. female reference), Day (session day), Tone (trial number), Handler2/3 (handler identity), Shock_hist (prior shock that day), and Weight (z-scored body weight), as well as the Sex:Day interaction. Predictors with consistent directionality across cohorts are shown in bold. The effect of Sex is highlighted. *p < 0.05, **p < 0.01, ***p < 0.001

**Figure 5.**
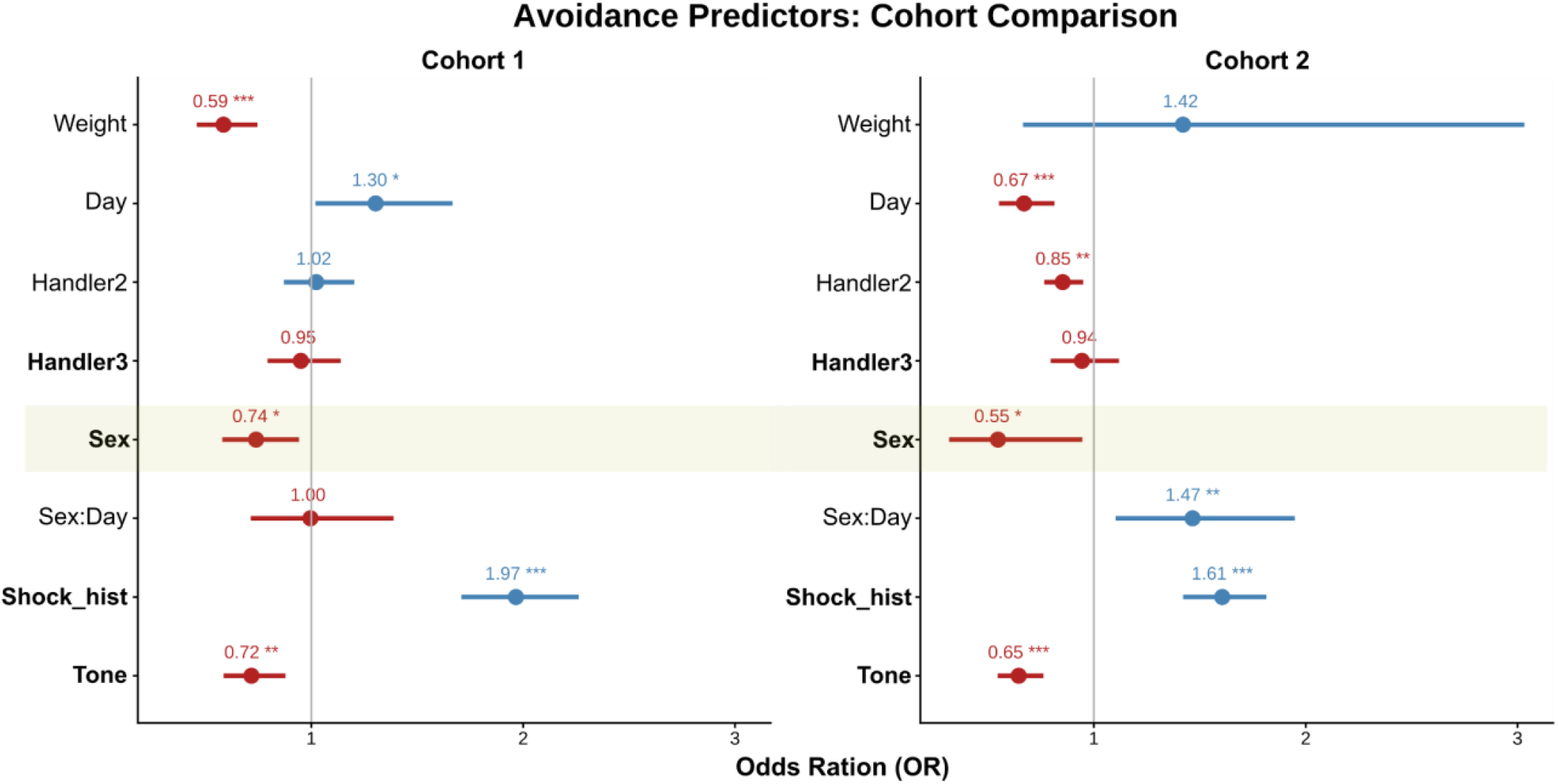
Comparison of predictors for avoidance behavior across cohorts. Forest plots display Odds Ratios (ORs) with 95% confidence intervals from the beta component of the avoidance GLMMs for Cohort 1 (left) and Cohort 2 (right). ORs greater than 1 (blue) are associated with more time on the platform, while values less than 1 (red) are associated with less. The vertical line at 1 represents no effect. Predictors with consistent directionality across cohorts are shown in bold. The effect of Sex is highlighted. *p < 0.05, **p < 0.01, ***p < 0.001.

**Figure 6.**
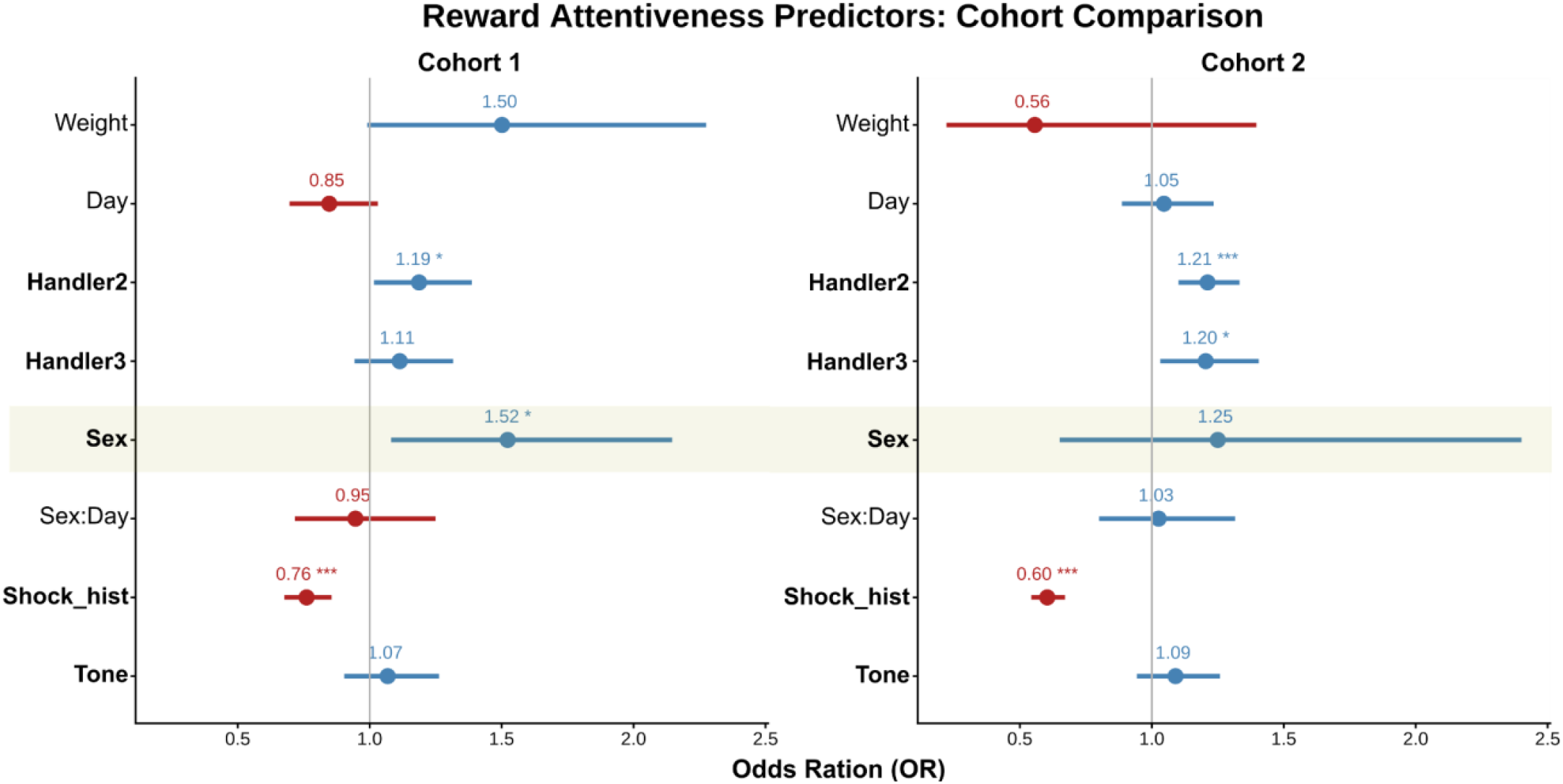
Comparison of predictors for reward attentiveness across cohorts. Forest plots display Odds Ratios (ORs) with 95% confidence intervals from the beta GLMMs for Cohort 1 (left) and Cohort 2 (right). ORs greater than 1 (blue) indicate a positive association with attentiveness, while values less than 1 (red) indicate a negative association. The vertical line at 1 represents no effect. Predictors with consistent directionality across cohorts are shown in bold. The effect of Sex is highlighted. *p < 0.05, **p < 0.01, ***p < 0.001.

### Estrous Cycle Analysis

To explore whether avoidance and approach strategies were modulated by hormonal state, we conducted a supplementary analysis on the combined female dataset (n=10) from the stable phase of behavior (days 20-25). Behavior was modeled as a function of a binary factor comparing the high-hormone phases (Proestrus/Estrus) against the low-hormone phases (Metestrus/Diestrus).

#### Reward-Seeking Behavior

In the ZINB model, estrous cycle phase (Proestrus/Estrus vs. Metestrus/Diestrus) was not a significant predictor of the number of bar presses (p = 0.129; Figure 7-A). Similarly, in the zero-inflation part of the model, the cycle phase did not predict the likelihood of an animal engaging reward-seeking entirely (p = 0.981). However, behavior remained sensitive to other variables; SH significantly reduced bar pressing (β = -0.25, z = -6.39, p < 0.001) and increased the probability of making zero presses (β = 0.71, z = 3.46, p < 0.001).

**Figure 7.**
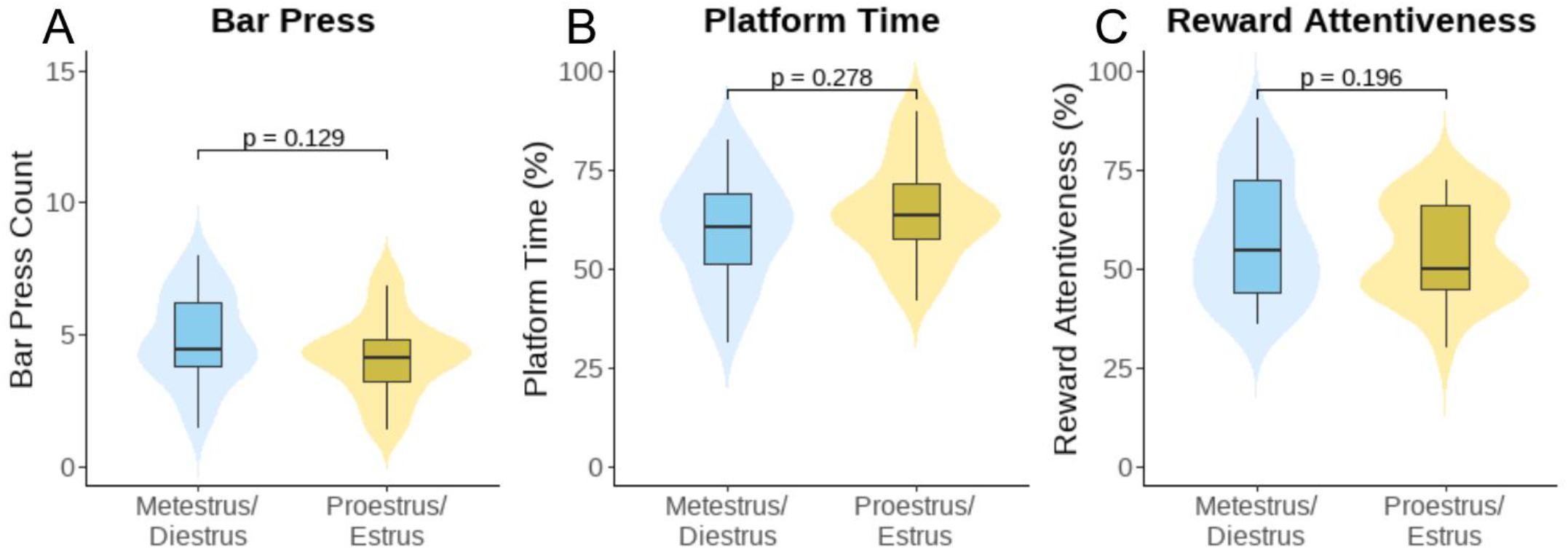
Estrous cycle does not impact stable-phase performance. Comparison of (A) Bar Press Count, (B) Platform Time (%), and (C) Reward Attentiveness (%) between low-hormone (Metestrus/Diestrus) and high-hormone (Proestrus/Estrus) states. Data are from the stable phase (Days 20-25) for all females (n=10, combined) and are visualized using violin plots with overlaid boxplots. P-values from the GLMM analysis are indicated.

#### Avoidance Behavior

The two-part hurdle-beta model for avoidance revealed no significant effect of estrous cycle phase. In the hurdle component, estrous phase did not predict the decision to remain on the platform (p = 0.991), which was instead positively associated with both day (β = 0.83, z = 2.82, p = 0.005) and trial number (β = 1.00, z = 2.90, p = 0.004). Similarly, in the beta component, estrous phase did not predict time spent avoiding (p = 0.278; Figure 7-B). This duration was strongly increased by SH (β = 0.62, z = 9.02, p < 0.001) and decreased by trial number (β = - 0.34, z = -3.58, p < 0.001).

#### Reward Attentiveness

In the beta GLMM for reward attentiveness, no significant main effect of estrous cycle phase was found (p = 0.196; Figure 7-C). Attentiveness was, however, significantly predicted by other factors. Both handling groups H2 (β = 0.24, z = 3.80, p < 0.001) and H3 (β = 0.31, z = 3.80, p < 0.001) showed increased attentiveness compared to the reference group. As in other models, SH was also an important predictor, significantly reducing attentiveness (β = -0.34, z = -5.79, p < 0.001).

## Discussion

Confirming our initial hypothesis, the present study demonstrates robust and replicable sex differences in resolving a high-stakes approach-avoidance conflict, with females consistently prioritizing safety while males exhibit more persistent reward-seeking. These findings provide strong evidence for sex-specific behavioral strategies that align with and extend the existing literature on dimorphic threat responses and risk-reward preferences. This divergence seems consistent with other tasks where females often adopt stable, risk-averse strategies, while males are more influenced by recent outcomes and less deterred by negative consequences (Grissom & Reyes, 2019; Chen et al., 2021a; Grissom et al., 2024). Critically, interpreting these results requires moving beyond a simple conclusion that females are “more responsive” to threats. Instead, as Shansky (2018) argues, these behavioral divergences likely represent qualitatively different, and potentially adaptive, strategies for navigating high-conflict situations.

Our results fit squarely within a broader framework of active versus passive defensive behaviors. Preclinical literature consistently shows that when faced with a threat, males often adopt passive, immobile responses like freezing, whereas females tend to engage in active, mobile strategies (Dalla & Shors, 2009, Gruene et al., 2015). This active strategy in females has been termed “anxioescapic behavior,” characterized by movement-based responses, such as darting or escape, rather than simple threat response potentiation (Shanazz et al., 2022, Bangasser & Cuarenta, 2021). The PMA task is exceptionally well-suited to capture this strategic divergence. By providing an unambiguous safe location, it allows for the expression of an active avoidance response that might be masked in paradigms that rely solely on freezing. The greater time females spent on the platform, therefore, likely reflects the expression of this active coping strategy rather than a simple inability to approach the reward.

The high-conflict nature of our paradigm, where rewards were only available under imminent threat, is crucial for revealing these sex-specific priorities. Rather than simply reflecting greater anxiety, this strategic divide may represent a sex difference in decision-making, as females in both human and animal studies show a strong tendency to avoid options associated with frequent negative outcomes (Van Den Bos et al., 2013; Grissom & Reyes, 2019). This strategic divide is not unique to PMA and has been observed in other approach-avoidance models. For instance, in the Vogel conflict test, where water-deprived rats are punished with a shock for drinking, females also display greater behavioral inhibition, accepting significantly fewer punished responses than males (Basso et al., 2011). This parallel finding reinforces the idea that females, across different types of conflict, adopt a more cautious, safety-prioritized strategy. Notably, Basso and colleagues (2011) also reported a pharmacological dissociation: while classic benzodiazepines were anxiolytic in both sexes, other compounds like SSRIs and buspirone only produced anxiolytic-like effects in males. This pharmacological dissociation provides an important indication: while some core circuits for processing conflict may be shared between sexes, the systems that modulate these circuits and ultimately shape the behavioral strategy seem sexually dimorphic. This suggests that the decision-making process involves both shared and sex-specific aspects of neurocircuitry and circuit functioning, which may be differentially modulated by neuromodulatory systems.

It is important to notice that, while behaviors like bar pressing and avoidance differed strongly between males and females, reward attentiveness showed no consistent sex differences. Both sexes appear to monitor the reward-associated cues, but females more readily suppress the motor-approach response in the face of threat. This aligns with the idea that males and females may adopt qualitatively different behavioral strategies (Shansky, 2018), where the female strategy involves a more cautious cost-benefit analysis that prioritizes safety despite persistent engagement with the prospect of reward. This dissociation between attention and action suggests the critical difference lies within the decision-making and threat-response regulating circuits that weigh reward against potential threat.

These distinct behavioral strategies likely arise from sexually dimorphic neural and endocrine mechanisms. The work of Bravo-Rivera et al. (2021), which characterized these behavioral phenotypes in rats, offers a potential neural basis for our findings. They demonstrated that an avoidance-preferring strategy, which we observed in females, was associated with heightened activity in the amygdala, a key hub for threat response. Furthermore, gonadal hormones may profoundly shape these circuits. This shaping occurs not only through the immediate influence of circulating hormones in adulthood (activational effects) but also through the permanent organizing of neural pathways during critical developmental periods (organizational effects) (McCarthy et al., 2012; 2015). The influence of high estrogen levels, for instance, on facilitating the extinction of conditioned fear would be an activational effect (Graham & Milad, 2014; Ramikie & Ressler, 2018). However, findings from Halcomb et al. (2024) in a modified PMA task revealed that persistent avoidance in female mice was significantly reduced by a glucocorticoid receptor (GR) antagonist administered during learning. This suggests that GR activation during the acquisition of the threat association, rather than activational effects of circulating gonadal hormones alone, may be a key mechanism that “defines” and maintains this avoidance strategy in females.

While our study did not find a significant modulation by the estrous cycle in our exploratory analysis, the role of hormonal fluctuations remains an important consideration (Li & Graham et. al, 2017). This is consistent with findings from both the modified PMA task (Halcomb et al., 2024) and the Vogel test (Basso et al., 2011), where the estrous cycle did not predict avoidance behavior. The influence of the estrous cycle can be, however, task-dependent, as Carvalho et al. (2021) found it impacted fear-potentiated startle but not conditioned freezing. Together, this suggests that while ovarian hormones are critical for the developmental organization of these sexually dimorphic circuits, the expression of a stable, learned behavioral strategy in a high-conflict task may be less sensitive to acute fluctuations than other measures of fear-like responses.

In conclusion, this study reveals that male and female rats employ fundamentally different strategies to resolve approach-avoidance conflict. Females adopt a safety-first strategy, consistent with a more active coping style that seems more resistant to change. Males, in contrast, employ a risk-prone, reward-oriented strategy. The translational importance of these findings is significant. The persistent avoidance shown by females mirrors a core clinical feature of anxiety and trauma-related disorders that is often a primary target for intervention. Understanding the distinct neurobiological pathways and mechanisms that support these strategies is a critical step toward developing more effective, personalized treatments. While acknowledging challenges in translating circuit-level findings across species (de Oliveira et al., 2021) the next critical step is to characterize how this circuit activity differs between sexes. This characterization may provide the necessary biomarkers for developing targeted neuromodulation therapies, such as deep brain stimulation, to ameliorate persistent avoidance.

## Acknowledgements

We acknowledge financial support from the US National Institutes of Health (R01MH123634), the Minnesota Medical Discovery Team on Addictions, and the MnDRIVE Brain Conditions Initiative. All opinions and data expressed herein are solely those of the authors and do not represent the position, interests, or opinions of any funding body.

